# An internal PDZ-binding motif in Densin-180 promotes activity-dependent SHANK scaffold remodelling

**DOI:** 10.64898/2026.07.07.736929

**Authors:** Yasumi Otani, Vignesh Srinivasan, Julia S. Töller, Till Kallem, Neil Ball, Igor Barsukov, Juha Saarikangas, Hans-Jürgen Kreienkamp, Benjamin T. Goult

**Affiliations:** Department of Biochemistry, Cell & Systems Biology, Institute of Systems, Molecular & Integrative Biology, University of Liverpool, Crown Street, Liverpool L69 7ZB, U.K; Helsinki Institute of Life Science, HiLIFE and 4 Faculty of Biological and Environmental Sciences, University of Helsinki, P.O. Box56, 00014, Helsinki, Finland; Institute of Human Genetics, University Medical Center Hamburg-Eppendorf, Hamburg, Germany; Liverpool Interdisciplinary Neuroscience Centre, University of Liverpool, Liverpool, L9 7AL, U.K

**Author notes:** Corresponding authors: Ben Goult, Hans-Jürgen Kreienkamp and Juha Saarikangas. These authors contributed equally to this work (Yasumi Otani, Vignesh Srinivasan, Julia S. Töller).

## Abstract

SHANK proteins form core postsynaptic density (PSD) scaffolds that organise signalling complexes through multiple interaction domains, including PDZ domains that typically recognise C-terminal peptide motifs. Here, we identify an internal PDZ recognition mechanism that links Densin-180 to SHANK and promotes SHANK scaffold assembly. We map SHANK binding to an internal PDZ-binding motif in Densin-180 (residues 843-863) and show by NMR spectroscopy and fluorescence polarisation that this motif binds SHANK1-3 PDZ domains with high affinity and specificity, competing with canonical C-terminal ligands. The crystal structure of the Densin-180-SHANK complex reveals that Phe858 inserts into the hydrophobic pocket of the PDZ domain despite the absence of a terminal carboxylate. In neurons, this interaction mediates Densin-180 recruitment to dendritic spines and drives activity-dependent reorganisation of postsynaptic SHANK3 assemblies, whereas mutation of Phe858 disrupts both processes. Disrupting the Densin-180-SHANK interaction using a Densin-180-derived peptide impairs activity-dependent spine structural plasticity, accompanied by defects in PSD organisation and actin cytoskeleton remodelling. These findings define an internal mode of PDZ recognition and reveal how Densin-180 couples neuronal activity to SHANK scaffold remodelling.

## Introduction

Synaptic scaffold proteins organise the molecular architecture that underlies signalling and plasticity at excitatory synapses. Densin-180 (encoded by the *LRRC7* gene) is a postsynaptic scaffold protein enriched in the PSD. Through multiple protein interaction domains, Densin-180 regulates dendritic spine structure and synaptic signalling pathways (Apperson et al., 1996; Carlisle et al., 2011; Walikonis et al., 2001). Genetic studies have linked loss-of-function and missense variants in *LRRC7* to neurological and neuropsychiatric disorders (Khan et al., 2026; Willim et al., 2024), highlighting the importance of understanding the molecular mechanisms by which Densin-180 organises synaptic signalling complexes.

Our previous studies (Quitsch et al., 2005) as well as others (Witte et al., 2019) have demonstrated that Densin-180 associates with SHANK (SH3 and multiple ankyrin repeat domains) in the rodent brain *in vivo* and suggested that this interaction may contribute to the organisation of postsynaptic scaffold complexes. However, the molecular mechanism and structural basis of the Densin-180/SHANK interaction have remained unclear.

The SHANK family comprises three members encoded by the *SHANK1*, *SHANK2* and *SHANK3* genes. SHANK proteins function as major structural hubs within the PSD, linking neurotransmitter receptors, signalling molecules and the actin cytoskeleton (Duffney et al., 2015; Jia et al., 2025; Kreienkamp, 2008; Liu et al., 2026). Each SHANK protein contains a conserved domain architecture including a Shank/ProSAP N-terminal (SPN) domain, a set of six ankyrin repeats, an SH3 domain, a class I PDZ (PSD-95/Discs-large/ZO-1) domain, a proline-rich region and a C-terminal SAM (sterile α motif) domain. These modular interaction domains, together with SAM-mediated multimerisation, enable SHANK proteins to assemble large signalling complexes that regulate synaptic architecture and dendritic spine morphology (Baron et al., 2006; Hassani Nia et al., 2024; Naisbitt et al., 1999; Sala et al., 2001). Consistent with this central scaffolding role, mutations in SHANK genes are strongly associated with autism spectrum disorders and other neurodevelopmental conditions (Leblond et al., 2014).

A key interaction module within SHANK proteins is the PDZ domain, which recruits binding partners to the postsynaptic scaffold (Hassani Nia et al., 2024; Naisbitt et al., 1999; Sala et al., 2001). PDZ domains are among the most abundant protein-protein interaction modules in synaptic scaffold proteins (Kim and Sheng, 2004). These domains typically recognise short peptide motifs located at the extreme C-terminus of their binding partners (Songyang et al., 1997). In canonical PDZ interactions, the terminal carboxylate and a hydrophobic residue of the ligand dock into a conserved groove within the PDZ domain. Based on the sequence of these motifs, PDZ ligands are commonly classified into several major classes, including the class I consensus sequence Ser/Thr-X-Φ-COOH (X; any amino acid, Φ; hydrophobic amino acid). This mode of recognition underlies many interactions that organise synaptic scaffold networks (Doyle et al., 1996; Im et al., 2003; Lee and Zheng, 2010).

However, alternative modes of PDZ recognition have occasionally been reported. In some cases, internal sequences can engage PDZ domains by adopting conformations that mimic canonical ligands, as shown, for example, for the SHANK1 PDZ domain (Ali et al., 2021; Li et al., 2026, 2024). The mechanisms by which such internal motifs interact with PDZ domains remain poorly understood. Furthermore, recognition of the SHANK PDZ domain via internal PDZ ligand motifs has so far not been described within the context of synaptic scaffolding in neuronal systems.

Against this background, the molecular mechanism by which Densin-180 engages SHANK proteins, and whether this interaction contributes to synaptic scaffold organisation, has remained unclear. Here we combine structural biology, biochemistry, cell biology and neuronal experiments to define how Densin-180 engages SHANK proteins and how this interaction promotes activity-dependent remodelling of postsynaptic scaffolds. Our findings establish an internal PDZ recognition mechanism that links Densin-180 to SHANK scaffolds, providing a molecular route for activity-dependent organisation of the PSD.

## Results

### The SHANK PDZ domain mediates the interaction with Densin-180

To identify the region of SHANK responsible for binding Densin-180, we generated a series of mRFP-tagged SHANK3 deletion constructs (Fig. 1A) and co-expressed them in HEK293T cells together with full-length GFP-tagged Densin-180. Densin-180 was immunoprecipitated from cell lysates using GFP-trap, and associated SHANK3 fragments were analysed by immunoblotting. SHANK3 fragments containing the PDZ domain (residues 576-668 in rat SHANK3; UniProt Q9JLU4-1) robustly co-immunoprecipitated with Densin-180 (Fig. 1B). In contrast, fragments comprising the N-terminal SPN, ankyrin repeat and SH3 domains were not efficiently co-precipitated with Densin-180. These results indicate that the PDZ domain of SHANK3 is necessary for the interaction with Densin-180.

**Figure 1.**
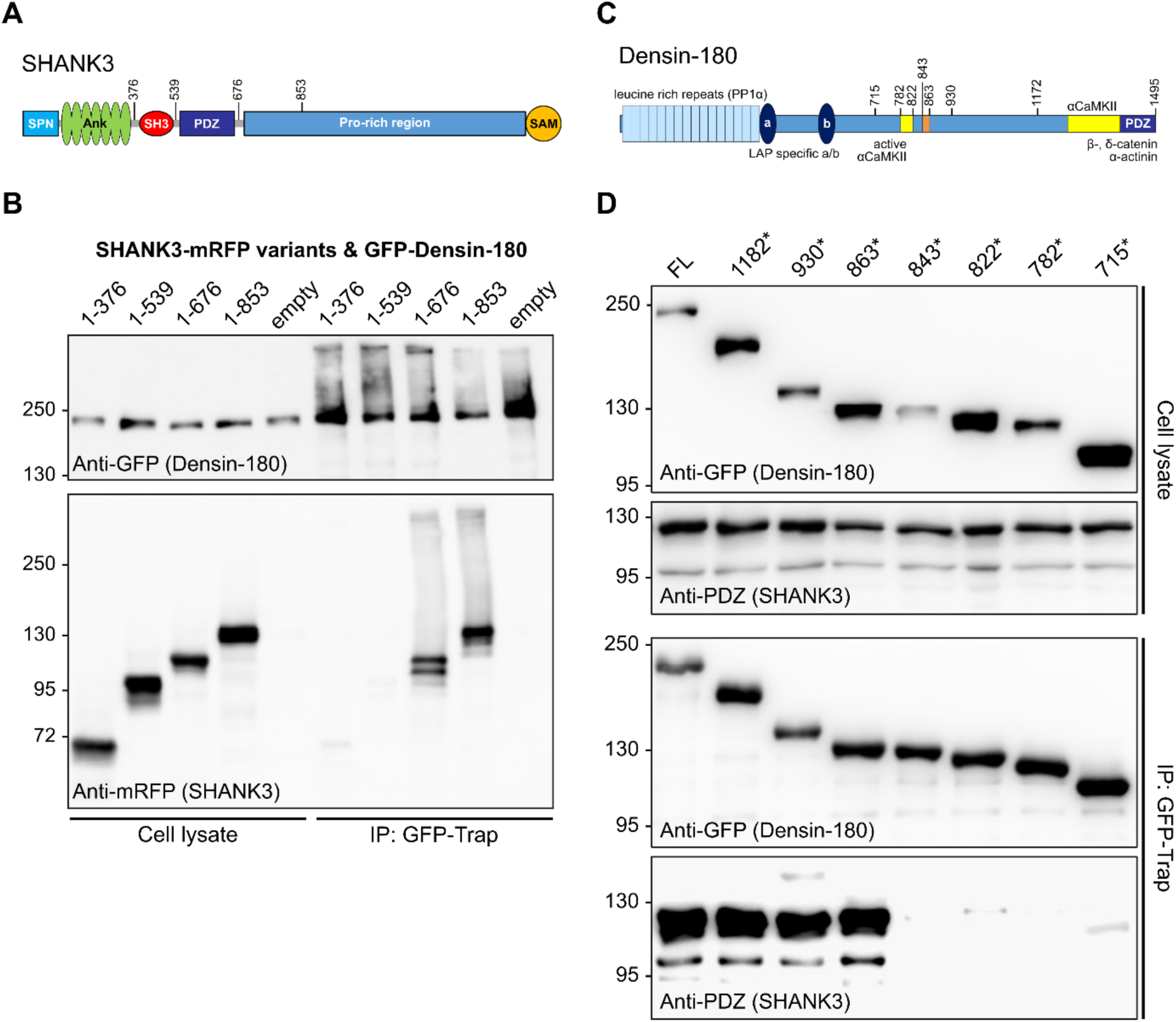
The SHANK PDZ domain binds an internal region of Densin-180. **(A)** Domain organisation of SHANK3 showing SPN, ankyrin repeat (Ank), SH3, PDZ, proline-rich and SAM domains. Boundaries of deletion constructs are indicated. (**B)** Co-immunoprecipitation assay mapping the SHANK3 region required for Densin-180 binding. mRFP-tagged SHANK3 fragments were co-expressed with GFP-tagged Densin-180 in HEK293T cells. Fragments containing the PDZ domain are efficiently co-immunoprecipitated, whereas N-terminal fragments lacking the PDZ domain show minimal interaction, indicating that the PDZ domain mediates binding. **(C)** Domain organisation of Densin-180 indicating truncation constructs used for mapping. **(D)** Co-immunoprecipitation assay mapping the SHANK-binding region in Densin-180. GFP-tagged truncation constructs were co-expressed with mRFP-tagged SHANK3 (residues 1-676). Fragments terminating at residue 863 retain binding, whereas truncation to residue 843 abolishes interaction, defining residues 843-863 (orange) as the minimal SHANK-binding region.

### An internal region of Densin-180 mediates SHANK binding

Densin-180 lacks a canonical C-terminal class I PDZ-binding motif, therefore the mechanism of its interaction with SHANK has remained unclear. To define the SHANK-binding region within Densin-180, we performed deletion mapping using GFP-tagged C-terminal truncations of Densin-180 (Fig. 1C,D). A minimal mRFP-tagged SHANK3 construct that retained binding (residues 1-676) was co-expressed in HEK293T cells with a series of GFP-tagged Densin-180 deletion constructs. Densin-180 variants were immunoprecipitated using GFP-trap, and associated SHANK3 was analysed by immunoblotting. Robust binding was observed with a Densin-180 fragment terminating at residue 863, whereas binding was abolished upon further truncation to residue 843 (Fig. 1D). These results localise SHANK binding to a short internal region spanning residues 843-863 (rat Densin-180; UniProt P70587-1; sequence QNWTRTPSPFEDRTAFPSKLE, Fig. 2A). Inspection of this sequence revealed a class I PDZ-binding consensus motif (Thr856-Ala857-Phe858). Notably, however, this motif is positioned internally rather than at the protein C-terminus.

**Figure 2.**
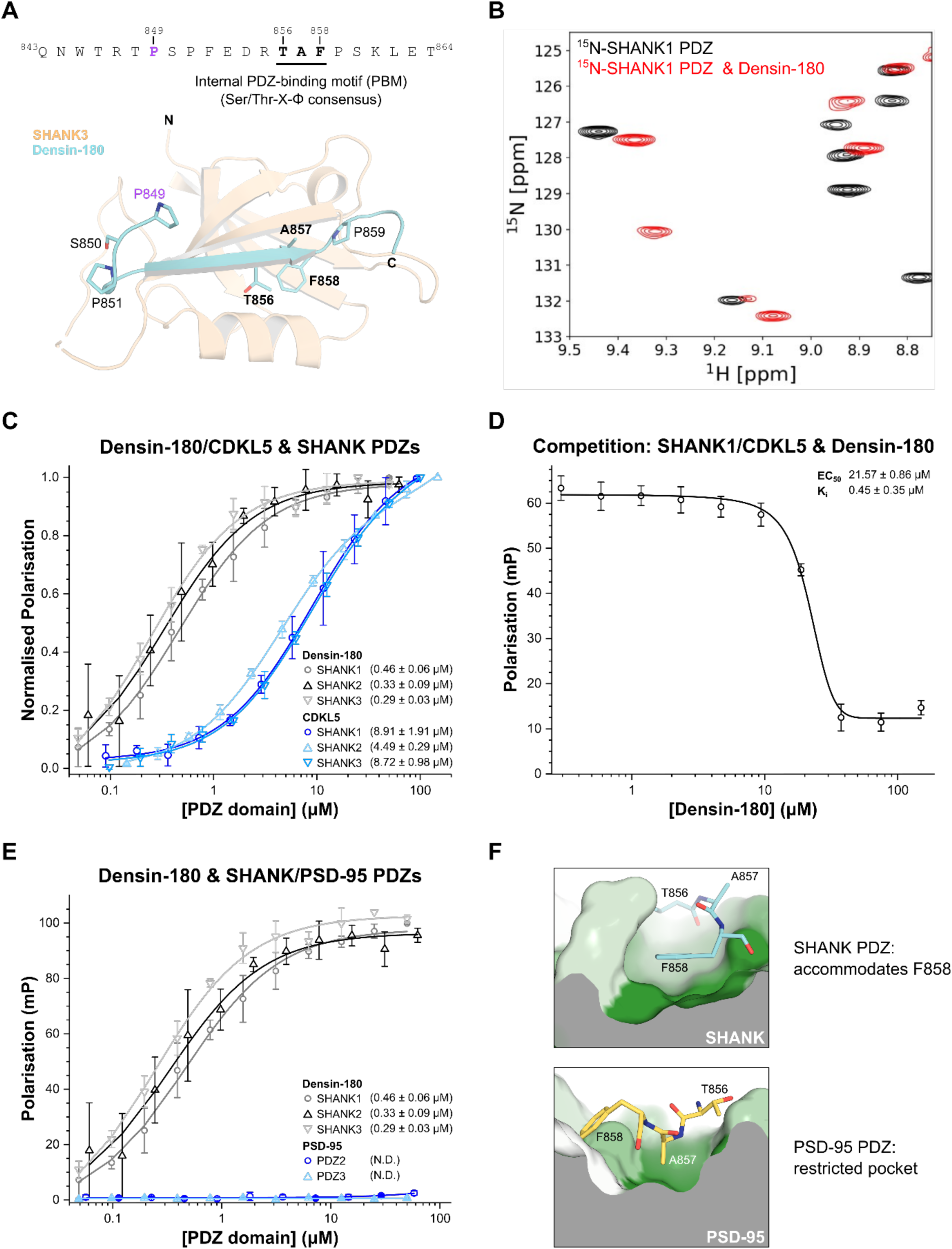
An internal PDZ-binding motif in Densin-180 engages SHANK PDZ domains. **(A)** Top, sequence of the internal Densin-180 SHANK-binding peptide (residues 843-864), with the class I PDZ-binding motif underlined. Bottom, crystal structure of the SHANK3 PDZ domain in complex with the Densin-180 peptide, showing insertion of the internal motif into the canonical PDZ ligand-binding groove. Key residues within and around the motif are indicated. **(B)** Overlay of ^15^N-HSQC spectra of SHANK1 PDZ domain alone (black) and after addition of Densin-180 peptide at a 1:1.2 molar ratio (red), demonstrating direct binding. **(C)** Fluorescence polarisation (FP) assay measuring binding of the Densin-180 peptide or the CDKL5 PDZ-binding peptide (residues 945-960) to SHANK PDZ domains. Peptides were fluorescently labelled via a non-native cysteine residue. Dissociation constants ± SE (µM) are indicated in the legend. All measurements were performed in triplicate. **(D)** Competition FP assay showing displacement of fluorescently labelled CDKL5 peptide from SHANK1 PDZ by unlabelled Densin-180 peptide, indicating binding to the canonical PDZ ligand-binding site. EC₅₀ and calculated K_i_ values are indicated. **(E)** FP assay measuring binding of the Densin-180 peptide to PDZ domains from SHANK proteins and PSD-95. The peptide binds all SHANK PDZ domains with comparable affinity but shows no detectable binding to PSD-95 PDZ domains, indicating SHANK-selective PDZ binding. ND, not determined. **(F)** Structural comparison showing that Phe858 is accommodated within the SHANK PDZ pocket, whereas the corresponding pocket in PSD-95 is more restricted. PDZ domains are coloured by hydrophobicity using the AAindex scale FASG890101 (Nakai et al., 1988), where green indicates hydrophobic residues and white indicates polar residues.

To test whether this sequence binds SHANK PDZ domains directly, we performed NMR chemical shift perturbation experiments using 15N-labelled SHANK1 PDZ. Addition of the Densin-180 peptide spanning residues 843-864 induced clear chemical shift perturbations in the HSQC spectrum, demonstrating direct binding in solution (Fig. 2B). We therefore next sought to define the structural basis of this interaction.

### Crystal structure reveals recognition of an internal PDZ-binding motif by the SHANK3 PDZ domain

To define how the internal Densin-180 motif engages SHANK PDZ domains, we determined the crystal structure of the SHANK3 PDZ domain bound to a Densin-180 peptide spanning residues 843-864. The structure was solved to 1.98 Å resolution, with data collection and refinement statistics reported in Table 1, and revealed that residues 856-858 of Densin-180 adopt the geometry of a class I PDZ ligand within the SHANK3 PDZ-binding groove (Fig. 2A). The bound peptide was well resolved in both the 2Fo-Fc electron density and simulated annealing Fo-Fc omit maps (Supplementary Fig. 1). Thr856 occupies the canonical Ser/Thr position, with its hydroxyl group contacting His718 in the SHANK3 PDZ domain, while Phe858 inserts into the conserved hydrophobic pocket normally occupied by the terminal hydrophobic residue of classical PDZ ligands. Thus, despite being located internally and lacking a free C-terminus, the Densin-180 motif preserves key recognition features of a canonical PDZ ligand.

**Table 1.** X-ray data collection and refinement statistics for SHANK3 PDZ (570-664) in complex with Densin-180 (843-864C).

|  |  |
| --- | --- |
| Synchrotron and Beamline | Diamond I24 |
| Space group | P2<sub>1</sub>2<sub>1</sub>2<sub>1</sub> |
| Molecule/ASU | 4 |
| Cell dimensions |  |
| *a*, *b*, *c* (Å) | 63.89, 71.34, 114.92 |
| $\alpha$, $\beta$, $\gamma$ (°) | 90, 90, 90 |
| Resolution (Å) | 47.61 - 1.98<br>(2.01 - 1.98)* |
| *I* / $\sigma$(*I*) | 7.6 (1.1) |
| CC(1/2) | 1.0 (0.4) |
| Completeness (%) | 99.9 (99.8) |
| Redundancy | 6.6 (5.7) |
| Refinement |  |
| Resolution (Å) | 47.64-1.98 |
| No. reflections | 37309 (1820) |
| *R*<sub>work</sub> / *R*<sub>free</sub> | 0.2010/0.2566 |
| No. atoms |  |
| Protein | 7308 |
| Water | 367 |
| *B*-factors (Å<sup>2</sup>) |  |
| Protein | 30.5 |
| Water | 44.5 |
| R.M.S. deviations |  |
| Bond lengths (Å) | 0.014 |
| Bond angles (°) | 2.16 |
| Ramachandran plot |  |
| Favoured/outlier (%) | 97/0.7 |
| Rotamer |  |
| Favoured/poor (%) | 92/8 |
| MolProbity scores |  |
| Clash score all atoms | 4.76 |
| PDB accession no. | 32CR |

Data collected from a single crystal.
\*Values in parentheses are for highest-resolution shell.

### Internal PDZ recognition defines SHANK-selective binding

Having established the structural basis of the interaction, we next quantified binding using fluorescence polarisation (FP) assays. Consistent with the crystal structure, the Densin-180 peptide bound to PDZ domains from SHANK1, SHANK2 and SHANK3 with comparable affinities (Fig. 2C), indicating that this interaction is conserved across the SHANK family. Notably, despite reproducing the canonical binding geometry observed in the crystal structure, the Densin-180 peptide bound with approximately ten-fold higher affinity than a canonical C-terminal PDZ-binding peptide derived from CDKL5 (Otani et al., 2025). Thus, high-affinity PDZ recognition is not restricted to C-terminal ligands.

The structure revealed that the Densin-180 peptide engages the canonical PDZ ligand-binding groove of SHANK. We therefore asked whether this internal ligand competes with canonical C-terminal PDZ ligands for the same binding site. To address this, we performed competition FP assays in which increasing concentrations of unlabelled Densin-180 peptide displaced a fluorescently labelled CDKL5 peptide from SHANK1 PDZ (Fig. 2D), indicating that both peptides bind to the same ligand-binding groove.

We next examined whether this interaction is specific to SHANK PDZ domains. PSD-95 is a major postsynaptic scaffold protein containing three class I PDZ domains. Although both SHANK and PSD-95 PDZ domains recognise class I PDZ-binding motifs, the Densin-180 peptide bound robustly to PDZ domains from all SHANK family members but showed no detectable binding to PSD-95 (Fig. 2E), demonstrating that the interaction is selective for SHANK PDZ domains despite the shared consensus sequence. Structural comparison provides a mechanistic explanation for this specificity. The Densin-180 motif contains phenylalanine 858, which inserts into the hydrophobic pocket of the SHANK PDZ domain, where it is accommodated by a relatively large binding cavity. In contrast, the corresponding pocket in PSD-95 PDZ domains is more restricted and is predicted to disfavour accommodation of this bulky hydrophobic residue, consistent with the lack of detectable binding (Fig. 2F).

Internal PDZ recognition is therefore not a generic property of PDZ domains but instead reflects specific structural compatibility with SHANK PDZ domains. Together, these data demonstrate that internal PDZ recognition can confer protein-level specificity beyond the canonical PDZ-binding consensus.

### Mutational analysis defines structural determinants of the internal PDZ-binding motif in Densin-180

Canonical PDZ interactions rely on insertion of a hydrophobic C-terminal residue into the PDZ binding pocket. The structure of the SHANK-Densin-180 complex revealed that Phe858 within the Densin-180 peptide occupies this same pocket despite the motif being located internally, adopting a geometry similar to that of canonical PDZ ligands (Figs. 2A and 2F). To test the importance of this residue, Phe858 was mutated to aspartate (F858D) to disrupt hydrophobic packing. FP experiments showed that the F858D mutation completely abolished binding to SHANK PDZ domains (Fig. 3A), demonstrating that this residue is essential for the interaction. Together with the ability of the wild-type Densin-180 peptide to compete with a canonical PDZ ligand for SHANK binding (Fig. 2D), these results define residues 856-858 as the core of the internal PDZ-binding motif.

**Figure 3.**
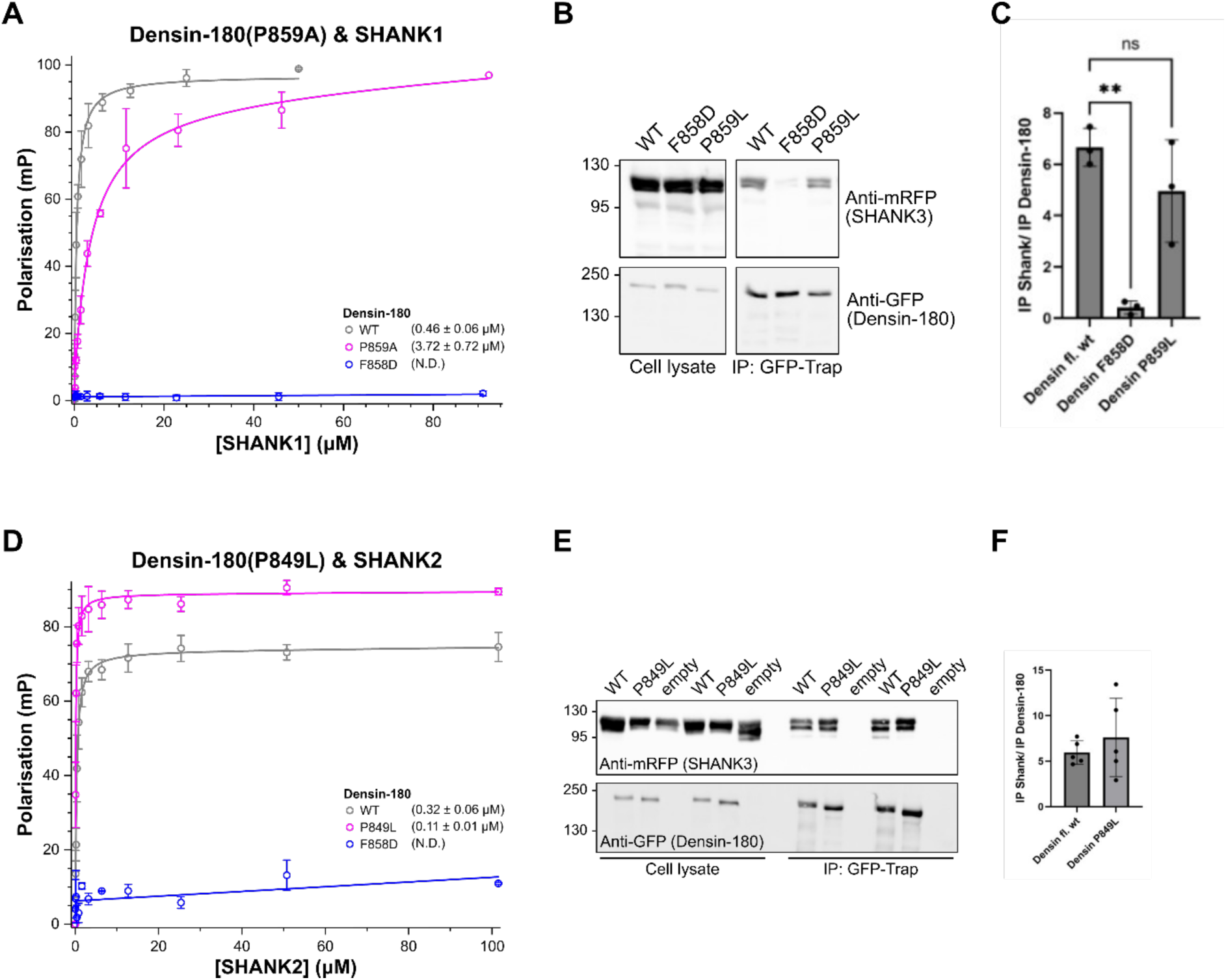
Mutational analysis defines structural determinants of the internal PDZ-binding motif. **(A)** Fluorescence polarisation (FP) assay measuring binding of Densin-180 wild-type (WT), F858D and P859A peptides to SHANK1 PDZ. Mutation of Phe858 abolishes binding, whereas P859A reduces affinity. Dissociation constants ± SE (µM) are indicated. **(B)** Co-immunoprecipitation assay of GFP-tagged Densin-180 WT, F858D, P859L and P849L variants co-expressed with mRFP-tagged SHANK3 (residues 1-676) in HEK293T cells. Densin-180 was immunoprecipitated using GFP-trap and co-precipitating SHANK3 was detected by immunoblotting. **(C)** Quantification of SHANK3 co-immunoprecipitation from (B). **(D)** FP assay measuring binding of Densin-180 wild-type and P849L peptides to SHANK2 PDZ, showing that the disease-associated flanking variant alters binding affinity. Dissociation constants ± SE are indicated. **(E)** Co-immunoprecipitation analysis of the P849L patient variant. **(F)** Quantification of SHANK3 co-immunoprecipitation for the P849L variant. Data represent mean ± SEM from independent experiments. FP measurements were performed in triplicate. ND, not determined. ns, not significant; **p≤0.01.

Immediately following the Thr856-Ala857-Phe858 (TAF) motif, Pro859 introduces a bend in the peptide backbone that positions Phe858 within the binding pocket. We therefore tested whether altering this adjacent proline affects SHANK binding. *In vitro*, substitution of Pro859 with alanine weakened binding by ∼8-fold (WT K_d_ 0.45±0.13 μM; P859A K_d_ 3.6±1.8 μM), indicating that this position contributes to high-affinity engagement (Fig. 3A).

We next examined whether residues flanking the motif modulate the interaction. A Pro849Leu (P849L) variant was recently detected in a patient with a neurodevelopmental disorder with phenotypic features (Khan et al., 2026) similar to those of other Densin-180-linked variants (Willim et al., 2024). Fluorescence polarisation measurements showed that this mutation alters binding affinity (Fig. 3D), indicating that residues surrounding the core motif influence PDZ engagement.

To determine whether these effects are recapitulated in cells, we co-expressed GFP-tagged Densin-180 variants with mRFP-tagged SHANK3 (residues 1-676) in HEK293T cells. Following immunoprecipitation of GFP-tagged Densin-180, associated SHANK3 fragments were analysed by immunoblotting. Consistent with the biochemical data, the F858D mutation abolished SHANK3 binding, confirming the requirement for a hydrophobic residue at this position (Fig. 3B,C). We also examined an independently generated full-length Densin-180 Pro859Leu variant. This variant retained detectable SHANK3 binding, indicating that substitution at Pro859 weakens but does not abolish the interaction in the context of full-length Densin-180 (Fig. 3B,C). In contrast, the P849L patient variant did not significantly alter SHANK3 binding in this assay (Fig. 3E,F), despite its modest effect on binding affinity measured by fluorescence polarisation.

Together, these results establish a structural framework in which Phe858 forms the core PDZ anchor, whereas neighbouring residues fine-tune binding affinity. We next examined how this interaction contributes to SHANK scaffold organisation in the functional context of neuronal synapses.

### The Densin-180 internal PDZ-binding motif regulates its recruitment to neuronal dendritic spines

We next assessed whether mutations within the internal PDZ-binding motif (F858D) or adjacent to it (P859L, P849L) impair Densin-180 recruitment to dendritic spines. Rat primary cortical neurons were co-transfected with GFP-tagged Densin-180 constructs together with mScarlet-C1 to delineate dendritic morphology, then fixed and imaged by confocal microscopy. To quantify spine localisation, we measured Densin-180-GFP fluorescence intensity in dendritic spines and adjacent dendritic shafts. Whereas wild-type Densin-180 showed comparable fluorescence intensity in spines and dendrites (spine/dendrite ratio: ∼1), all three mutants were preferentially localised to dendritic shafts (spine/dendrite ratio ∼0.6), indicating impaired spine recruitment (Fig. 4A,B). These results indicate that the internal PDZ-binding motif and its adjacent residues contribute to Densin-180 localisation at dendritic spines.

**Figure 4.**
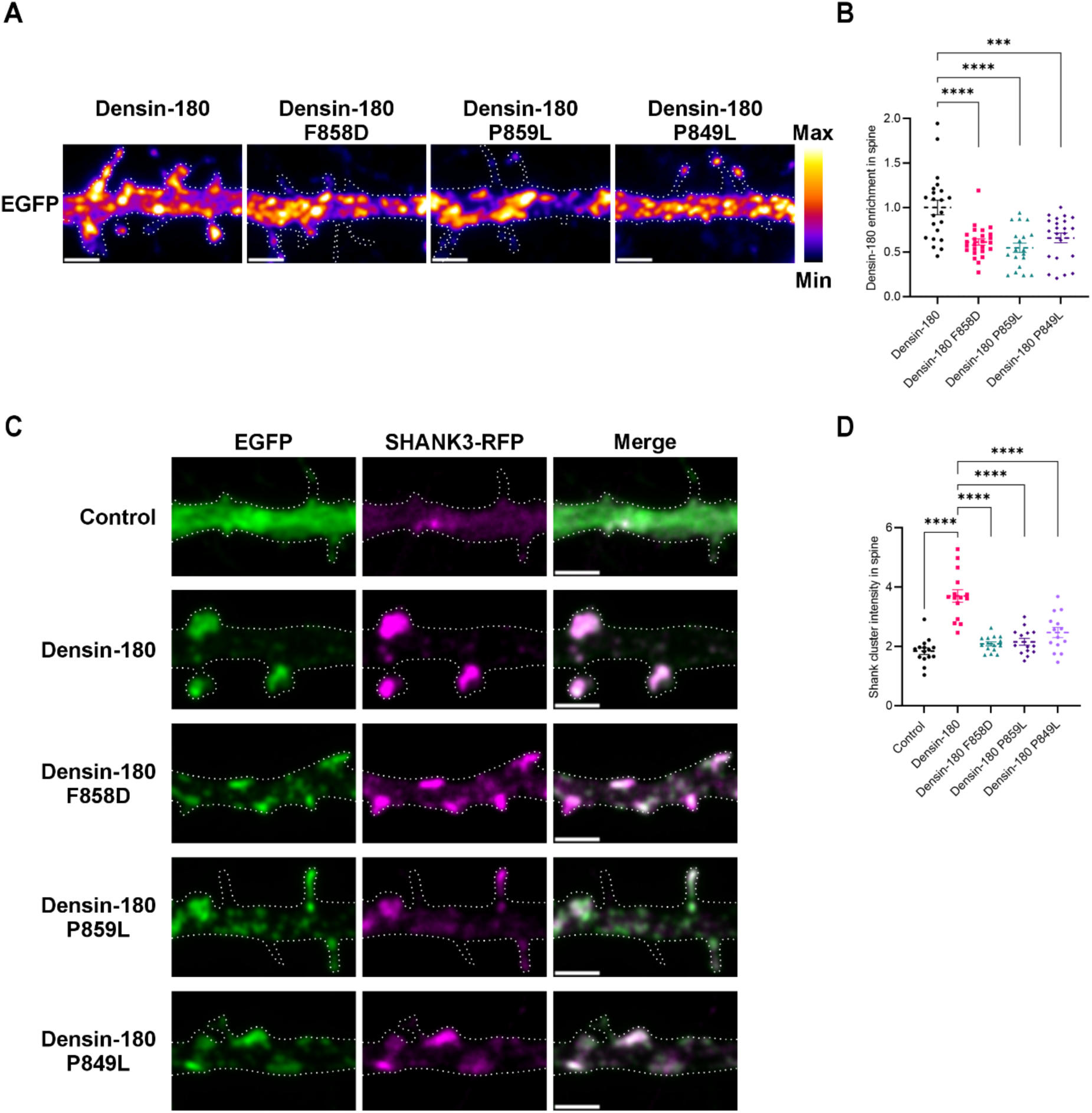
The internal PDZ-binding motif regulates Densin-180 localisation and SHANK3 clustering in dendritic spines. **(A)** Representative images of DIV14 primary cortical neurons expressing GFP-tagged Densin-180 WT, F858D, P859L or P849L. GFP signal is shown using the Flame LUT. Right, Flame LUT intensity scale. **(B)** Quantification of Densin-180 spine enrichment, calculated as mean GFP intensity in dendritic spines normalised to the adjacent dendritic shaft. **(C)** Representative images showing the effect of Densin-180 WT and mutants on SHANK3 clustering in dendritic spines. Neurons expressing EGFP alone were used as controls. Densin-180 constructs are shown in green, SHANK3-mRFP in magenta and merged signal in grey. **(D)** Quantification of SHANK3 cluster enrichment in spines, calculated as SHANK3 puncta mean intensity normalised to total SHANK3 mean intensity in the dendritic shaft. (B,D) Each data point represents an individual dendritic shaft containing 10-30 spines from 8-10 neurons per condition. Mean spine values from the same shaft were averaged. Statistical analysis was performed using ordinary one-way ANOVA with multiple comparisons. ***p≤0.001, ****p≤0.0001. (A,C) White dotted lines indicate dendritic morphology boundaries. Scale bars, 2 µm.

### The Densin-180-SHANK PDZ interaction controls SHANK scaffold assembly in neurons

Having established that the internal PDZ-binding motif contributes to Densin-180 recruitment to dendritic spines, we next asked whether this interaction promotes SHANK scaffold assembly. We co-expressed GFP-tagged WT or mutant Densin-180 together with mRFP-tagged SHANK3 and BFP-CAAX as a morphology marker in primary cortical neurons. After 24 h of expression, neurons were fixed and imaged by confocal microscopy, and SHANK3 clustering within dendritic spines was quantified (Fig. 4C,D).

In control neurons expressing GFP alone, SHANK3 formed discrete puncta within dendritic shafts and adjacent spines (Fig. 4C, first row). Expression of WT Densin-180 significantly increased SHANK3 cluster formation within dendritic spines (Fig. 4C, second row; Fig. 4D), indicating that Densin-180 promotes postsynaptic SHANK scaffold assembly.

We next examined whether this function depends on the Densin-180-SHANK PDZ interaction. Mutation of the core PDZ anchor, Phe858Asp (F858D), markedly impaired the ability of Densin-180 to promote SHANK3 clustering (Fig. 4C, third row; Fig. 4D). A similar reduction was observed with the neighbouring P859L mutation, indicating that perturbing this motif compromises SHANK scaffold assembly (Fig. 4C, fourth row; Fig. 4D).

Finally, we examined the disease-associated Pro849Leu (P849L) variant, which lies immediately upstream of the motif. Although this variant produced only modest effects on SHANK binding *in vitro*, it significantly reduced SHANK3 cluster formation in dendritic spines compared with WT Densin-180 (Fig. 4C, fifth row; Fig. 4D), demonstrating that subtle perturbations of the internal PDZ-binding motif can have pronounced effects on postsynaptic scaffold assembly.

Together, these data show that Densin-180 promotes postsynaptic SHANK scaffold assembly through its internal PDZ-binding motif and indicate that disruption of this mechanism may contribute to neurodevelopmental disease.

### The PDZ-binding motif of Densin-180 is required for activity-dependent SHANK3 and PSD-95 assembly at synapses

Structural reorganisation and expansion of the PSD scaffold, including recruitment of SHANK3, is a hallmark of activity-dependent synaptic strengthening (Bosch et al., 2014; Droogers and MacGillavry, 2023; Clavet-Fournier et al., 2024). We examined the role of the Densin-180-SHANK interaction in this process by expressing a GFP-tagged Densin-180 peptide containing the SHANK-binding motif (residues 843-864), designed to disrupt endogenous Densin-180-SHANK interactions through competition. We then monitored endogenous SHANK3 and PSD-95 organisation following chemically induced LTP (cLTP), triggered by glycine-mediated activation of NMDARs, using immunofluorescence and BFP-CAAX as a morphology marker.

As expected, cLTP induced significant accumulation of endogenous SHANK3 and PSD-95 in dendritic spines of control EGFP-expressing neurons (Fig. 5A, first and second rows, Fig. 5B-D). In contrast, expression of EGFP-Densin-180(843-864) strongly impaired activity-dependent recruitment of SHANK3 and PSD-95 (Fig. 5A, third and fourth rows, Fig. 5B-D). Quantification of SHANK3 cluster intensity also revealed reduced basal SHANK3 accumulation in dendritic spines upon disruption of the Densin-180-SHANK interaction (Fig. 5C). Collectively, these data support a role for Densin-180 in promoting SHANK3 clustering in spines and indicate that the Densin-180-SHANK interaction is required for activity-dependent remodelling of the PSD during synaptic strengthening.

**Figure 5.**
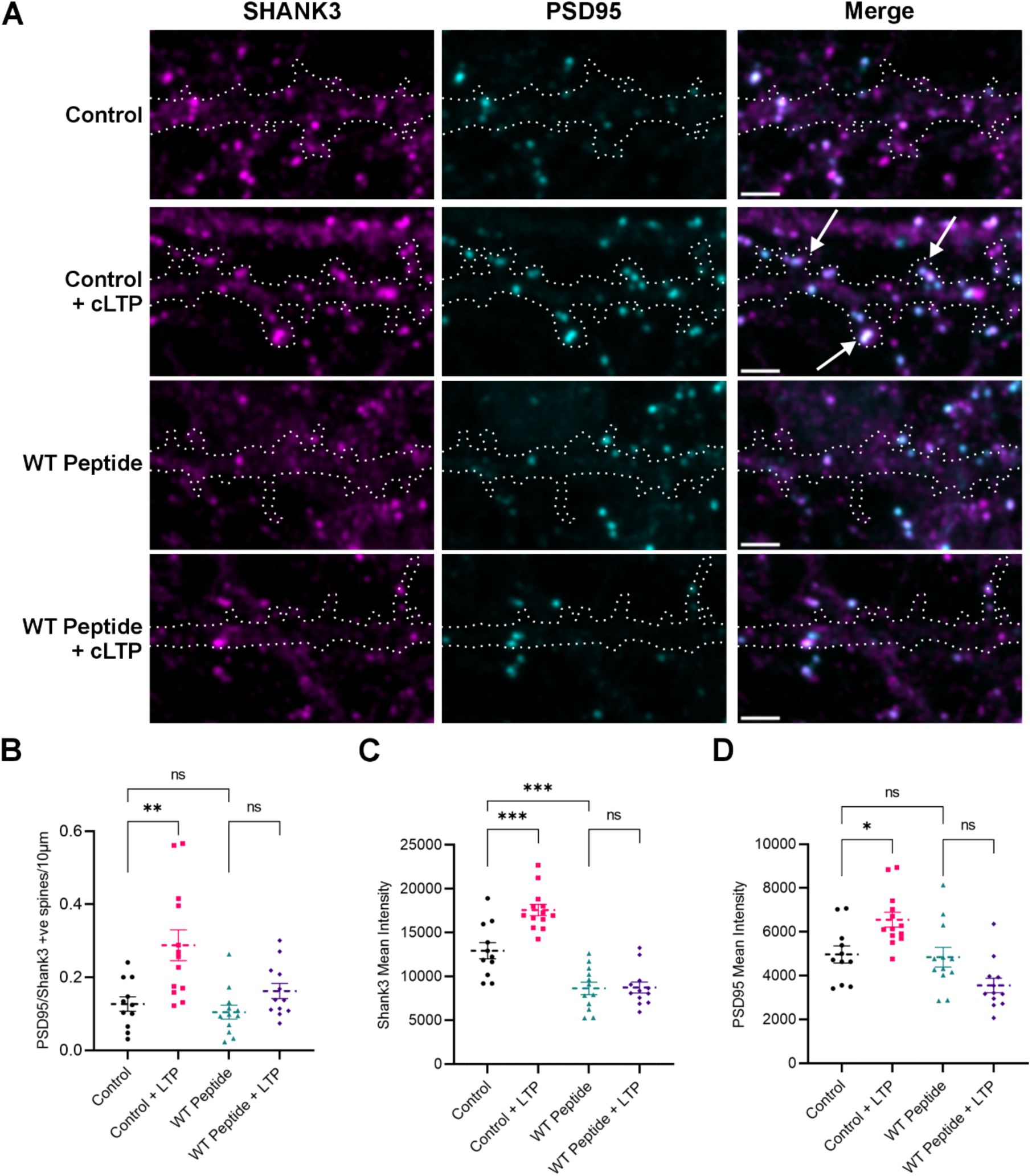
The Densin-180-SHANK interaction is required for activity-dependent SHANK3 and PSD-95 assembly at synapses. **(A)** Representative images of DIV14 primary cortical neurons expressing EGFP alone or EGFP-tagged Densin-180(843-864), before or after 30 min glycine-induced cLTP. Endogenous SHANK3 is shown in magenta and PSD-95 in cyan. White arrows indicate SHANK3/PSD-95 double-positive spines. **(B)** Quantification of SHANK3/PSD-95 double-positive spines per 10 µm of dendritic shaft. **(C)** Quantification of SHANK3 cluster mean intensity in SHANK3/PSD-95 double-positive spines. **(D)** Quantification of PSD-95 cluster mean intensity in SHANK3/PSD-95 double-positive spines. (B-D) Each data point represents an individual dendritic shaft from 8-10 neurons per condition. Statistical analysis was performed using ordinary one-way ANOVA with multiple comparisons. *p≤0.05, **p≤0.01, ***p≤0.001, ns, not significant. (A) Representative dendritic shaft regions are shown. White dotted lines outline dendritic boundaries. Scale bars, 2 µm.

### The Densin-180-SHANK interaction is required for activity-dependent structural plasticity of dendritic spines

We next assessed whether the Densin-180-SHANK interaction is required for activity-dependent structural plasticity by quantifying spine morphology in neurons expressing either EGFP or EGFP-tagged Densin-180(843-864), with or without cLTP induction. mScarlet was used as a morphological marker. Spine morphology was classified as mushroom, thin or stubby, and the distribution of spine types was determined for each condition (Fig. 6C). As expected, cLTP increased the proportion of mature mushroom spines in control EGFP-expressing neurons (Fig. 6A,D). In contrast, this increase was abolished in neurons expressing the Densin-180 peptide (Fig. 6B,D). No significant differences in spine morphology were observed between control and peptide-expressing neurons under basal conditions. These findings indicate that the Densin-180-SHANK interaction is required for activity-dependent maturation of dendritic spines (Fig. 6C,D).

**Figure 6.**
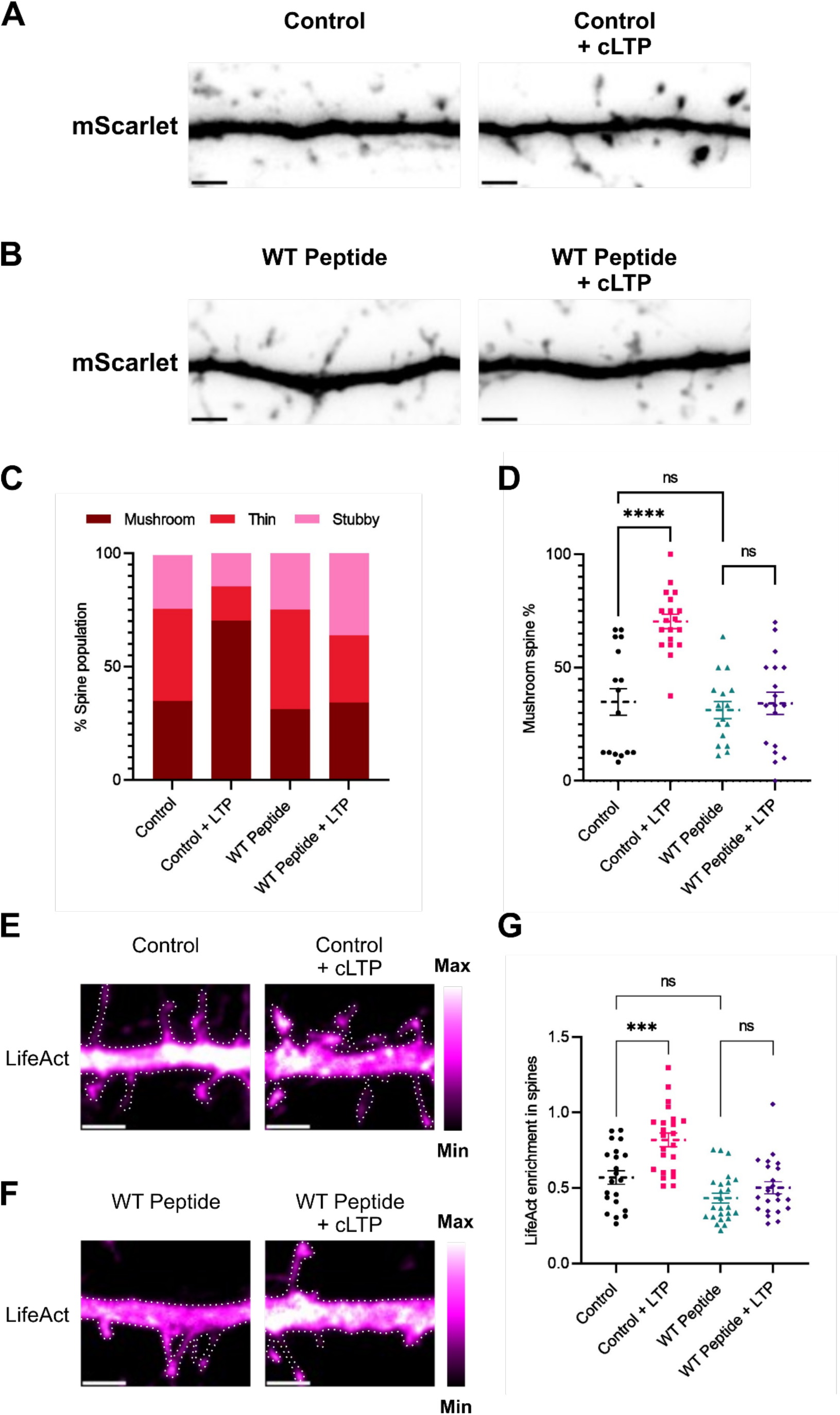
The Densin-180-SHANK interaction is required for activity-dependent structural plasticity and F-actin enrichment in dendritic spines. **(A-B)** Representative images of dendritic spines in neurons expressing **(A)** EGFP alone or **(B)** EGFP-tagged Densin-180(843-864), with or without cLTP. mScarlet is shown using an inverted greyscale LUT. **(C)** Distribution of mushroom, thin and stubby spine types, expressed as a percentage of total spines per dendritic shaft. **(D)** Quantification of mushroom spine percentage per dendritic shaft. **(E-F)** Representative images of mRFP-tagged LifeAct in neurons expressing **(E)** EGFP alone or **(F)** EGFP-tagged Densin-180(843-864), with or without cLTP. LifeAct is shown using the Magenta Hot LUT. Right, Magenta Hot LUT intensity scale. **(G)** Quantification of LifeAct spine enrichment, calculated as mean LifeAct intensity in spines normalised to the adjacent dendritic shaft. (D, G) Each data point represents an individual dendritic shaft from 8-10 neurons for each condition. Statistical analysis was performed using ordinary one-way ANOVA with multiple comparisons, *** p ≤ 0.001, **** p ≤ 0.0001, ns, not significant. (A-B, E-F) Representative dendritic shaft regions are shown. White dotted lines outline dendritic boundaries. Scale bars, 2 µm.

Structural plasticity of dendritic spines depends on reorganisation of the actin cytoskeleton and SHANK proteins are important regulators of this process (MacGillavry et al., 2016; Sarowar and Grabrucker, 2016). To determine whether the defects in structural plasticity were associated with impaired F-actin assembly, we quantified F-actin in spines following cLTP using mRFP-tagged LifeAct in neurons expressing EGFP alone or EGFP-Densin-180(843-864), with BFP-CAAX serving as a morphology marker. As expected, cLTP increased F-actin accumulation in dendritic spines of control neurons (Fig. 6E,G). In contrast, expression of the Densin-180 peptide markedly impaired activity-dependent F-actin accumulation (Fig. 6F,G), indicating that disruption of the Densin-180-SHANK interaction compromises activity-dependent actin remodelling during spine structural plasticity (Fig. 6G).

Together, these findings identify the Densin-180-SHANK interaction as a molecular mechanism that couples neuronal activity to SHANK scaffold remodelling and dendritic spine maturation.

## Discussion

In this study, we identify a mechanism that promotes synaptic plasticity by coupling Densin-180 to activity-dependent assembly of SHANK scaffolds. We show that a short internal sequence within Densin-180 directly engages the canonical ligand-binding groove of SHANK PDZ domains despite lacking the free C-terminal carboxylate that typically defines PDZ ligands. The crystal structure of the Densin-180-SHANK complex and mutational analysis show that phenylalanine 858 inserts into the canonical hydrophobic pocket of the SHANK PDZ domain, while neighbouring residues shape the conformation of the internal motif and influence binding affinity. We demonstrate that this interaction is important for SHANK3 assembly in spines, contributing to activity-dependent remodelling of the PSD and structural plasticity of dendritic spines. Together, these findings reveal an unexpected mode of PDZ recognition that links molecular binding specificity to the assembly of postsynaptic scaffolds.

PDZ domains are best known for recognising short peptide motifs located at the extreme C-terminus of their binding partners, where the terminal carboxylate and a hydrophobic residue occupy a conserved binding groove within the PDZ domain. This mode of recognition underlies many synaptic scaffold interactions and has led to the view that PDZ-mediated signalling networks are largely organised through C-terminal ligands. The interaction described here represents a distinct mode of PDZ engagement in which an internal sequence within Densin-180 preserves the binding geometry of a canonical PDZ ligand despite lacking a free C-terminus. These findings indicate that PDZ domains can accommodate internal motifs that reproduce key features of canonical ligands, expanding the potential repertoire of PDZ-mediated interactions within synaptic scaffolds. The SHANK PDZ domain can therefore accommodate both conventional C-terminal ligands, such as CDKL5 (Otani et al., 2025), and internal ligands exemplified by Densin-180. Together with the ability of these ligands to compete for the same PDZ-binding surface, this suggests that SHANK PDZ domains function as competitive interaction hubs whose composition can be remodelled during synaptic activity.

Although the crystal structure reveals that the Densin-180 motif reproduces the key recognition features of a canonical class I PDZ ligand, the interaction displays substantially higher affinity than the canonical C-terminal CDKL5 peptide. This suggests that faithful engagement of the canonical PDZ-binding pocket is not, by itself, sufficient to define binding affinity. Instead, additional contacts outside the core PDZ-binding motif, or conformational features unique to internal ligands, may further stabilise the complex. In the structure, Thr856 occupies the canonical Ser/Thr position and contacts His718 in the SHANK3 PDZ domain, while Phe858 inserts into the hydrophobic pocket used by the terminal residue of classical PDZ ligands. This interaction appears to be facilitated by local peptide conformation, as Pro859 introduces a bend in the backbone that helps position Phe858 within the binding pocket. Consistent with this, mutation of Phe858 abolishes PDZ binding, while substitution of Pro859 weakens the interaction, indicating that both residues contribute to stabilising the internal PDZ ligand conformation.

Rather than acting solely as a constitutive postsynaptic scaffold, our data support a model in which Densin-180 becomes dynamically engaged during synaptic activity, where its internal PDZ-binding motif promotes assembly of SHANK-containing postsynaptic scaffolds. Densin-180 is enriched in postsynaptic densities (Apperson et al., 1996; Quitsch et al., 2005) and the interaction between Shank and Densin-180 in the rodent brain *in vivo* has been confirmed in several studies (Quitsch et al., 2005; Witte et al., 2019). Our data show that the PDZ-binding motif contributes both to Densin-180 localisation within dendritic spines and to SHANK scaffold assembly, indicating a reciprocal interaction that supports PSD organisation. Consistent with this model, expression of full-length Densin-180 enhanced SHANK3 clustering in dendritic spines, whereas mutations that disrupted the internal PDZ-binding motif, as well as expression of a competitive peptide, impaired SHANK scaffold assembly. Activity-dependent recruitment of SHANK3 during cLTP likewise required an intact Densin-180-SHANK interaction, and disruption of this interaction impaired PSD organisation, dendritic spine maturation and actin cytoskeleton remodelling. This model is consistent with previous reports that neurons from Densin-180 knockout mice exhibit abnormal spine morphology and impaired synaptic plasticity (Carlisle et al., 2011; Wang et al., 2017). Our results provide a mechanistic explanation for these phenotypes by identifying Densin-180 as a regulator of activity-dependent SHANK scaffold assembly. More broadly, these findings suggest that Densin-180 functions not simply as a constituent of the PSD, but as an active organiser whose recruitment promotes higher-order assembly of SHANK-containing scaffold complexes during synaptic plasticity.

Patients carrying mutations in either *SHANK* or *LRRC7*/Densin-180 share a molecular aetiology characterised by disrupted PSD architecture and defects in synaptic plasticity, accompanied by overlapping neurological phenotypes (Wan et al., 2022; Willim et al., 2024). In this context, we examined the role of a recently identified *LRRC7* missense variant linked to neurodevelopmental disorder (Khan et al., 2026), which affects a residue immediately upstream of the PDZ-binding motif (Pro849). We show that expression of the P849L variant impaired the ability of Densin-180 to promote SHANK clustering in neuronal dendritic spines, indicating that disruption of this region alters postsynaptic scaffold assembly. The more pronounced effect of P849L in neurons than in biochemical assays suggests that the cooperative environment of the PSD amplifies subtle perturbations of the Densin-180-SHANK interface into pronounced defects in scaffold assembly, providing a potential explanation for how relatively modest molecular changes can contribute to neurodevelopmental disease. Collectively, our results provide a molecular bridge that helps relate Densin-180- and SHANK-associated diseases to one another at the level of synaptic scaffold organisation. More broadly, our work expands the principles governing PDZ-mediated interactions and identifies the Densin-180-SHANK interaction as a molecular mechanism linking internal PDZ recognition to activity-dependent reorganisation of synaptic architecture.

## Methods

### Cloning

The native and mutant PDZ-binding motif of Densin-180 (residues 843-864) were ordered as synthetic genes (GeneArt, Life Technologies). These regions were sub-cloned into both an EGFP-tagged mammalian expression vector (EGFP-C1 with a modified multiple cloning site (MCS) based on pET47b) and a bacterial expression vector encoding SUMO as a solubility tag (based on pET47b with a region encoding SUMO inserted between the His-tag and HRV-3C protease site). Subcloning was performed using ligation independent cloning (LIC) where the Densin-180 region was inserted between the XmaI and SacI sites in the MCS of the linearised vectors with the addition of a non-native C-terminal cysteine (encoded in the reverse primer) to permit maleimide-fluorophore labelling.

Expression constructs coding for rat Lrrc7/Densin-180 (NM_001393674.1, full length) in pEGFP-N2, leading to a C-terminal fusion with EGFP, as well as in pEGFP-C1, leading to N-terminal GFP, have been described (4, 6). For expression in neurons, the pHAGE vector carrying the Lrrc7-EGFP sequence was used (4). Vectors coding for rat SHANK3 in C-terminal fusion with mRFP in the pmRFP-N backbone have been described (Hassani Nia et al., 2020).

Missense variants were introduced by site directed mutagenesis using the Quik-Change kit from Agilent, following the manufacturer’s instructions; for each mutant, two complementary oligonucleotides (Sigma-Aldrich, Steinheim, Germany) carrying the desired variant (F858D, P859L, S850D, S850A, P849L) were used in the reaction. Mutations were introduced both in pEGFP-N2 and pHAGE-based vectors. For deletion mapping, stop codons were introduced in the pEGFPC-1 based vector at desired positions. All constructs were verified by Sanger sequencing along the Lrrc7 coding sequence. All DNA constructs used in this study are available from the corresponding authors upon request.

### Protein expression and purification

All PDZ domains from SHANK and PSD-95 were expressed and purified as His-tagged recombinant proteins. Codon-optimised synthetic genes (GeneArt, Life Technologies) encoding human PSD-95 (UniProt: P78352) PDZ2 (residues 155-249) and PDZ3 (residues 302-402), mouse SHANK1 PDZ (UniProt: D3YZU1, residues 654-762), mouse SHANK2 PDZ (UniProt: Q80Z38, residues 247-340), and mouse SHANK3 PDZ (UniProt: Q4ACU6, residues 570-664).

For protein expression, *E. coli* BL21(DE3) Star cells were transformed with the appropriate plasmid and grown in LB medium supplemented with 100 µg/mL ampicillin or 50 µg/mL kanamycin. Cultures (750 mL LB) were inoculated with 7 mL starter culture and grown at 37 °C until OD_600_ reached 0.7-0.8. Protein expression was induced with 0.4 mM IPTG and cultures were incubated overnight at 20 °C.

For production of ^15^N-labelled proteins for NMR experiments, cells were grown in M9 minimal medium containing 1 g/L ^15^N-ammonium chloride as the sole nitrogen source.

For SUMO-Densin-180 peptide production, BL21(DE3) Star cells co-expressing chaperones from the pG-KJE8 plasmid (Takara Bioscience) were used.

Cells were harvested by centrifugation at 6000 × g for 10 min and resuspended in nickel affinity chromatography buffer A (20 mM Tris-HCl pH 8.0, 500 mM NaCl, 20 mM imidazole) or resuspension buffer (50 mM Tris-HCl pH 8.0, 250 mM NaCl, 10% (w/v) glycerol, 10 mM MgCl₂).

Cells were lysed by sonication and clarified lysates were applied to a 5 mL HisTrap HP nickel affinity column (Cytiva). Proteins were eluted and dialysed overnight into 20 mM Tris-HCl pH 8.0, 50 mM NaCl. Where required, the His-tag was removed using AcTEV protease (Invitrogen). Further purification was performed by anion exchange chromatography using a 5 mL HiTrap Q HP column (Cytiva).

For FP and NMR experiments, SUMO tags were removed by HRV-3C protease digestion following IEX purification. Purified proteins were buffer-exchanged into PBS (10 mM Na₂HPO₄, 1.8 mM KH₂PO₄, pH 7.4, 137 mM NaCl, 2.7 mM KCl) for fluorescence polarisation assays or into NMR buffer (12 mM NaH₂PO₄, 6 mM Na₂HPO₄, pH 6.5, 50 mM NaCl) for NMR experiments.

### Fluorescence Polarisation Assay (FP)

Peptides were either synthesised by GLBiochem (Shanghai) or expressed and purified recombinantly.

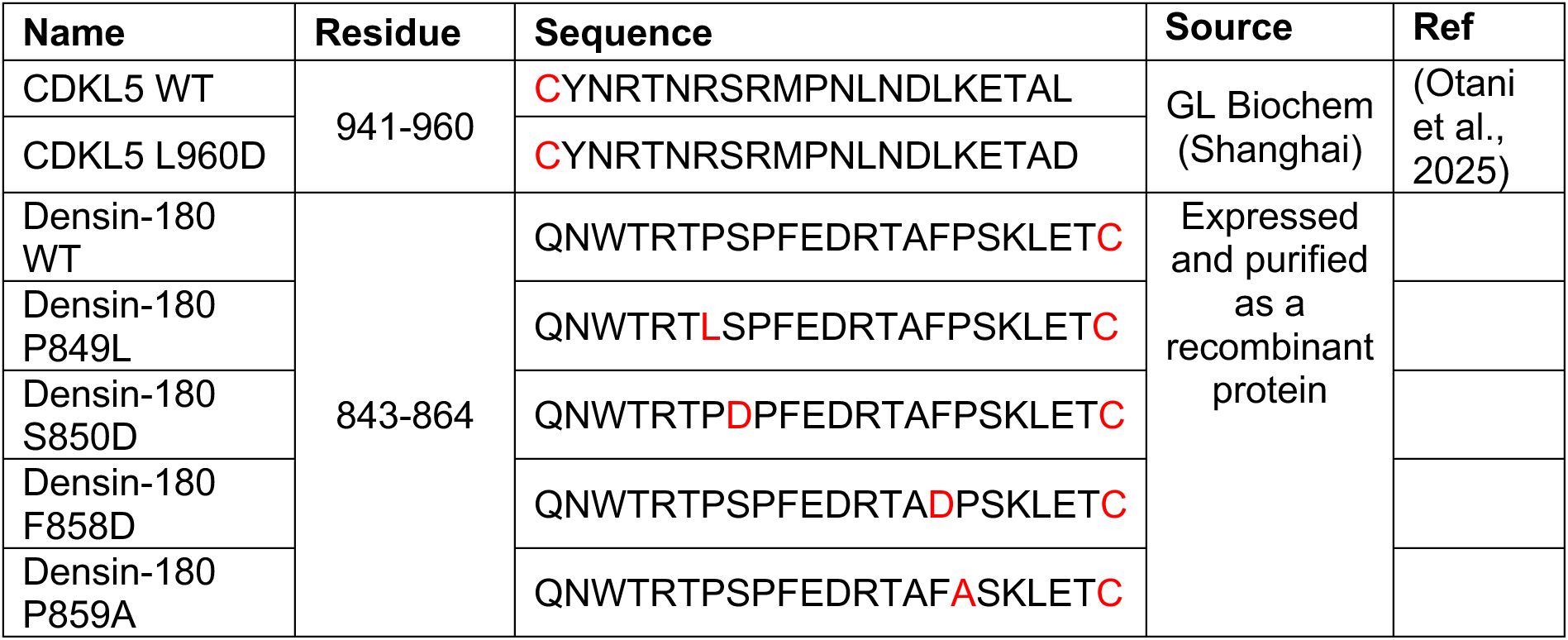

Peptides contained a non-native cysteine at either the N-terminus (CDKL5 peptides) or C-terminus (Densin-180 peptides) to enable conjugation to maleimide-fluorescein (Thermo Fisher Scientific). Peptides were labelled according to the manufacturer’s instructions.

Binding assays were performed by titrating purified PDZ domains from SHANK or PSD-95 against fluorescently labelled peptides in PBS. Proteins were prepared as twofold serial dilutions and mixed with labelled peptide in assay buffer. Samples were incubated for 15 min at room temperature to allow the binding reaction to reach equilibrium prior to measurement.

Fluorescence polarisation was measured at 25 °C using a HIDEX Sense microplate reader (excitation 485 ± 10 nm; emission 535 ± 20 nm). Binding curves were fitted using a one-site total binding model in OriginPro (Origin Lab). All measurements were performed in triplicate.

### Competition fluorescence polarisation assay

For competition assays, SHANK1 PDZ domain was incubated with fluorescently labelled CDKL5 C-terminal peptide to form a complex in PBS. Unlabelled SUMO-Densin-180 peptide was prepared as a twofold serial dilution and titrated into the SHANK1 PDZ-CDKL5 peptide complex. Fluorescence polarisation was measured at 25 °C using a HIDEX Sense microplate reader (excitation 485 ± 10 nm; emission 535 ± 20 nm). The EC_50_ value was determined by fitting the data to a Hill equation and the K_i_ was calculated using the following equation (using a K_d_ for the CDKL5-SHANK interaction determined using a regular FP assay):

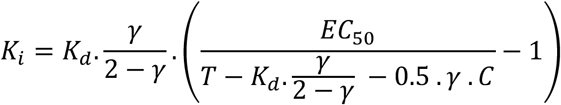

where T is the total SHANK concentration, C is the total concentration of fluorescent CDKL5 competitor and:

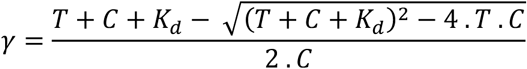

### Nuclear Magnetic Resonance (NMR)

^15^N-labelled SHANK PDZ domains were prepared at a final concentration of 150 µM in NMR buffer supplemented with 2 mM DTT and 5% (v/v) D₂O. Densin-180 peptides were added in the same buffer to the indicated molar ratios. ^1^H-^15^N HSQC spectra were collected at 298 K on a Bruker Avance III 700 MHz spectrometer equipped with a cryoprobe. Spectra were processed using TopSpin (Bruker) and analysed using CCPN Analysis (Skinner et al., 2015).

### X-ray crystallography

X-ray crystallography was used to determine the structure of the SHANK3 PDZ domain (570-664) in complex with Densin-180(843-864). The N-terminal His tag was cleaved by TEV protease at a 1:40 molar ratio overnight at 4 °C in 20 mM Tris pH 8.0 and 50 mM NaCl. TEV protease and the cleaved His tag were removed using a HiTrap Q HP anion exchange column (Cytiva). The cleaved protein was dialysed into 20 mM Tris pH 8.0 and 150 mM NaCl, concentrated to 15 mg/mL, flash-frozen in liquid nitrogen and stored at −80°C.

The SHANK3 PDZ domain was mixed with synthetic Densin-180(843-864C) peptide at a 1:1.1 molar ratio to a final concentration of 10 mg/mL in 20 mM Tris pH 8.0, 150 mM NaCl and 1 mM DTT. Sitting-drop vapour diffusion crystallisation drops were prepared by mixing protein solution and mother liquor in a 1:1 ratio, with a total drop volume of 1 μL, over a 90 μL equilibration reservoir. The mother liquor contained 60 mM MgCl_2_•6H_2_O, 60 mM CaCl_2_•2H_2_O, 100 mM Tris/Bicine pH 8.5, 12% v/v PEG 500 MME and 6% (w/v) PEG 20000. Crystals were harvested and flash-frozen in mother liquor using liquid nitrogen.

X-ray diffraction data were collected remotely on the I24 beamline at Diamond Light Source, UK, using a DECTRIS EIGER2 9M detector at a wavelength of 0.62 Å. The xia2 3dii pipeline was used to index, integrate and scale the diffraction data. The structure was solved by molecular replacement with MOLREP (Vagin and Teplyakov, 2010), using an AlphaFold model of SHANK3 PDZ in complex with Densin-180(843-864C) as the search model. Iterative rounds of real-space and reciprocal-space refinement were performed using Coot (Emsley et al., 2010) and REFMAC (Murshudov et al., 2011) within the CCP4i2 suite (Winn et al., 2011). Data collection and refinement statistics are reported in Table 1. The structure has been deposited in the PDB with accession code 32CR. Final structural figures were prepared using PyMOL (Schrodinger, LLC).

### Cell culture and transient transfection

293T human embryonic kidney cells from ATCC were maintained in Dulbecco’s modified Eagle’s medium containing 10% foetal bovine serum and 1% penicillin/streptomycin. Cells were transiently transfected with 3 μg of each plasmid and 18 μL TurboFect Transfection Reagent (Thermo Scientific) in 950 μL serum-free DMEM according to the manufacturer’s instructions. Cells were regularly monitored for absence of mycoplasma.

### Immunoprecipitation from transfected cells

Transfected cells were lysed in immunoprecipitation (IP) buffer (50 mM Tris-HCl, pH 8, 120 mM NaCl, 0.5% NP40, 1 mM EDTA), followed by centrifugation at 20,000 × g for 15 min at 4 °C. GFP-tagged proteins from supernatants were immunoprecipitated using 20 μL of magnetic GFP-trap beads (ChromoTek, Munich, Germany) and incubated on beads for 1 hr. After washing, precipitates and input samples were processed for Western blotting. The following primary antibodies were used: mouse anti-GFP (Covance MMS-118P-500, RRID: AB_291290; WB: 1:3000); rat anti-RFP (ChromoTek 5F8, WB: 1:1000); rabbit anti SHANK PDZ (1:2000) (ref. Zitzer et al., 1999; PMID: 10551867).

### SDS-PAGE and western blot

Proteins were separated by SDS-PAGE (8%) and blotted on nitrocellulose membrane using a MINI PROTEAN II system (Bio-Rad). After blocking membranes with 5% milk powder/ TBS-T, primary antibodies in milk/TBS-T were incubated overnight at 4 °C. After washing in TBS-T, HRP-linked secondary antibodies were incubated at room temperature for 1 h. Membranes were washed and then processed for chemiluminescence; imaging was done using a ChemiDoc TM MP Imaging System (Bio-Rad). Images were processed and further analysed using Image Lab Software (Bio-Rad).

### Statistical analysis

Statistical comparisons and graphing of neuronal imaging datasets were performed using GraphPad Prism v.10.2.3 (GraphPad Software, San Diego, CA, USA).

### Primary cortical neuron culture and transfections

Rat cortical neuronal cultures were prepared as described previously (Sahu et al., 2019) with slight modifications. Time-mated Wistar females were acquired from Charles River Laboratories (Germany) and maintained at The Laboratory Animal Centre (LAC) of the University of Helsinki. All animal work was done following institutional guidelines.

Briefly, cortices were dissected from E18-E19 rat embryos and placed in ice-cold HBSS. Tissues were enzymatically dissociated to obtain the neurons, which were plated on glass coverslips coated with 0.1 mg/mL poly-L-lysine (PLL) (Sigma-Aldrich) and cultured in MACS Neuro Medium (Miltenyi Biotec) supplemented with MACS NeuroBrew®-21 (Miltenyi Biotec), L-Glutamine (Invitrogen), and penicillin-streptomycin (Gibco). At DIV7 and DIV10, half of the cell-culture medium was replaced with fresh MACS medium with supplements. All neurons were transfected on DIV13 and fixed 24 h later at DIV14 or subjected to glycine-induced cLTP prior to fixation.

At DIV13, 100,000 cortical neurons per coverslip were transfected with 2 µL of Lipofectamine 2000 (ThermoFisher) and 1.2 µg of DNA following the manufacturer’s recommendation. The following plasmids were used for transfections: pVITRO1-BFP-CAAX (Addgene, #184459), pmScarlet_C1 (Addgene, #85042), pEGFP-N1 and LifeAct-pRFP (a gift from Pirta Hotulainen, University of Helsinki), pHage-based Densin-180 constructs (generated in this study), EGFP-tagged PDZ-binding motif of Densin-180 WT (residues 843-864) (generated in this study), SHANK3-mRFP (Hassani Nia et al., 2020).

### Glycine-induced chemical long-term potentiation (cLTP)

cLTP was induced in primary cortical neurons as described before (Hlushchenko and Hotulainen, 2019). Neurons at DIV14 were incubated in bath solution containing 120 mM NaCl, 5 mM KCl, 1.3 mM CaCl2, 10 mM HEPES, 10 mM D-glucose, 0.5 μM tetrodotoxin citrate, 50 μM picrotoxin and 1 μM strychnine with pH adjusted to 7.33 with NaOH. After 10 min, bath solution was replaced with fresh bath solution supplemented with 200 μM glycine (cLTP stimulation medium) and incubated for 4 min. Control (no cLTP) neurons were treated with fresh bath solution without glycine and incubated for 4 min. Following this, solutions in both control and cLTP samples were replaced with fresh bath solution, and neurons were placed in the incubator for 30 min before fixation.

### Primary cortical neuron imaging

DIV14 neurons were fixed either directly or after cLTP induction with 4% paraformaldehyde (PFA) for 15 min at room temperature, washed three times with 1x phosphate-buffered saline (PBS), and mounted onto glass slides with ProLong Diamond Antifade (ThermoFisher Scientific).

For SHANK3 and PSD-95 immunostaining, fixed neurons on coverslips were washed with PBS, followed by permeabilisation and blocking with PBS containing 0.1% Triton-X (PBS-T) supplemented with 10% normal goat serum (Abcam, #ab7481) for 30 min at room temperature (RT) with shaking. Coverslips were then incubated with a rabbit anti-SHANK3 antibody (Synaptic Systems, Cat. No. 162 302, 1:600) and single-domain antibodies (sdAbs) against PSD-95 raised in camelid and conjugated to AlexaFluor647 (Synaptic Systems, Cat. No. N3702-AF647-L, 1:500) diluted in PBS-T containing 3% normal goat serum, for 2 hr at RT with gentle shaking. Coverslips were then washed with PBS 3x, 30 min, and incubated with goat anti-rabbit IgG (H+L) Secondary Antibody, Alexa Fluor 568 (Invitrogen #A-11011, 1:500), for 60 min at RT with gentle shaking. Finally, coverslips were washed with PBS 3X, 30 min, dipped in MQ and mounted onto glass slides with ProLong Diamond Antifade (ThermoFisher Scientific).

Samples were imaged using Olympus FV4000 confocal microscope (Light Microscopy Unit, HiLIFE, University of Helsinki) equipped with a UPLXAPO 60x/1.42 oil-immersion objective and SilVIR detectors. For imaging, 405 nm, 488 nm, 561 nm, and 640 nm solid-state lasers were used, with the pinhole set to 1 Airy unit. The images were acquired in 16-bit format at a resolution of 512×512 or 1024×1024. Images were further processed and analysed using Fiji ImageJ.

### Confocal image processing and analysis

Quantification of Densin-180 and LifeAct enrichment in dendritic spines relative to adjacent shaft was performed as follows with ImageJ (Yang et al., 2018). SUM intensity projections of Z-stacks were made, and regions of interest (ROIs) were manually drawn around the spine head boundary based on the morphology marker. Mean intensity values of Densin-180 or LifeAct were acquired for each spine. Mean intensity values of Densin-180 or LifeAct from the adjacent shaft were obtained by manually defining the ROI around the shaft boundary. The mean intensity values of Densin-180 or LifeAct for each spine were normalised to the adjacent shaft mean intensity from which the spine protrudes. The values of the dendritic spines from the same shaft were averaged and represented as single data-point for each shaft, which was then used for statistical analysis.

SHANK3 signal enrichment in clusters localised to synapses was assessed using a thresholding strategy in ImageJ. Maximum intensity projections (MIPs) were generated from the Z-stack images of all channels. BFP-CAAX morphology marker channel was thresholded using the Li algorithm to create a binary dendrite mask that also includes spine protrusions. Fill holes or close were applied to produce a continuous mask of the dendritic compartment. For SHANK3 granule detection, the MIP projection of the SHANK3 channel was thresholded with RenyiEntropy algorithm to create binary cluster masks. The thresholding was chosen to exclude diffuse SHANK3 background signals while capturing discrete clusters. The same threshold was applied across all images, which were all acquired with identical confocal imaging settings. The morphology mask was applied to the cluster mask to filter out the clusters within the given morphological boundary. Individual puncta were quantified using the Analyze Particles function with a minimum size filter to exclude noise artifacts. Mean SHANK3 intensity values of detected clusters were obtained by redirecting particle measurements to the original, unthresholded SUM intensity projection (SIP) of the image. A selection was generated from the morphological mask and applied to SHANK3 unthresholded SIP image to acquire the mean SHANK3 intensity of the entire dendritic shaft. SHANK3 enrichment into clusters was calculated as: Mean intensity in clusters / Mean intensity in entire dendritic shaft, providing a value of how much SHANK3 is concentrated in clusters relative to the diffuse background signal. Clusters localised within the spines were manually annotated and categorised as SHANK3 enrichment in clusters within each spine. The spine localised cluster mean intensity values from the same shaft were averaged and used for statistical analysis.

Fluorescent analysis of dendritic spine-localised SHANK3 and PSD-95 cluster intensities were performed with ImageJ. The thresholding protocol (described above) was applied with slight modifications to acquire dendrite morphology mask (from the BFP-CAAX and EGFP channels) and binary cluster mask of PSD-95 granules from the MIP projection of respective channels. The morphology mask was applied to the PSD-95 cluster mask to detect only the clusters located within the morphology. PSD-95 cluster positive spines that are also exhibited SHANK3 cluster signals were manually annotated and counted. Density of spines double positive for SHANK3/PSD-95 clusters was calculated by normalising the number of double-positive spines to the total number of spines in each analysed shaft. These values were expressed as number of double positive spines per 10 µm length of shaft. To obtain the cluster intensity values, individual puncta were quantified using the Analyze Particles function with a minimum size filter to exclude noise artifacts. PSD-95 and SHANK3 intensities of spine-localised clusters (mean intensity values) were obtained by redirecting particle measurements to the original, unthresholded SUM intensity project (SIP) of the SHANK3 and PSD-95 channel images. PSD-95 and SHANK3 cluster intensities from the same shaft were averaged and represented as a single data-point for each shaft. The analysis parameters and thresholding strategy were kept identical across all samples, and all samples were imaged with the same microscopy settings.

Dendritic spine morphology was quantified manually using Fiji/ImageJ. Individual spines were identified along a continuous dendritic shaft segment and measured independently for each shaft. The following morphometric parameters were obtained from each spine:

- Total spine length: Distance from the base of the spine to the distal edge of the spine head.
- Spine head width: Widest point of the spine head.
- Spine neck width: Narrowest point of the spine neck.

Only spines with discernible protrusion from the dendritic shaft and not obstructed by adjacent structures were included for analysis. Spine classification was performed by morphometric approach described earlier, based on relative spine dimensions such as head:neck ratio and length:head ratio (Swanger et al., 2011; Schünemann et al., 2025).

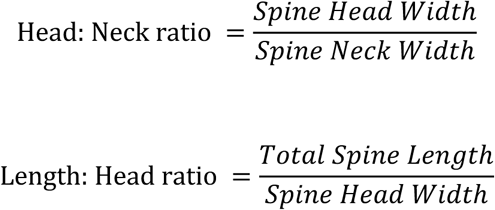

Spine classification based on the following ratios was applied consistently across all samples.

- Mushroom spines: Head: Neck ratio ≥ 1.5
- Thin Spines: Head: Neck Ratio ≤ 1.5 and Length:Head ratio > 2
- Stubby spines: Head: Neck Ratio ≤ 1.5 and Length:Head ratio ≤ 2

The proportion of mushroom, thin and stubby spines was calculated for each dendritic shaft and expressed as a percentage of the total number of analysed spines within that shaft. Each dendritic shaft segment served as a biological unit for the purpose of statistical analysis. The analysis was performed across multiple dendritic shaft segments from different neurons in each condition. For each spine sub-type, percentage values from different shafts were averaged and depicted as stacked bar graph to show the population distribution across the spine sub-type for each sample. Mushroom spine population percentage was plotted as a plot to represent changes in mushroom spine population statistics.

For all quantifications, images of 10-15 shafts from 8-10 neurons obtained from independent primary neuron isolations were used.

## Acknowledgements

B.G. was supported by a Cancer Research UK Programme Grant (CRUK-A21671) and a British Heart Foundation Special Project Grant (SP/F/23/150045). V.S. and J.S. were supported by the Magnus Ehrnrooth Foundation (to V.S.), Research Council of Finland (363743), and Jane and Aatos Erkko Foundation (240046). J.T. was supported by a grant from Werner-Otto-Stiftung. The NMR spectra were measured at the LIV-SRF High-Field NMR Facility, University of Liverpool. We thank Diamond Light Source, UK, for synchrotron access (MX15591 BAG). We acknowledge the services and facilities of the Laboratory Animal Centre (LAC) of the University of Helsinki and the Light Microscopy Unit, Institute of Biotechnology, supported by HiLIFE and Biocenter Finland.

## Data accessibility

The atomic coordinates and structure factors for the SHANK3 PDZ domain in complex with the Densin-180 peptide have been deposited in the Protein Data Bank under accession code 32CR. All other data supporting the findings of this study are included within the article and its supplementary information.

## Supplementary Figure

**Supplementary Figure 1.**
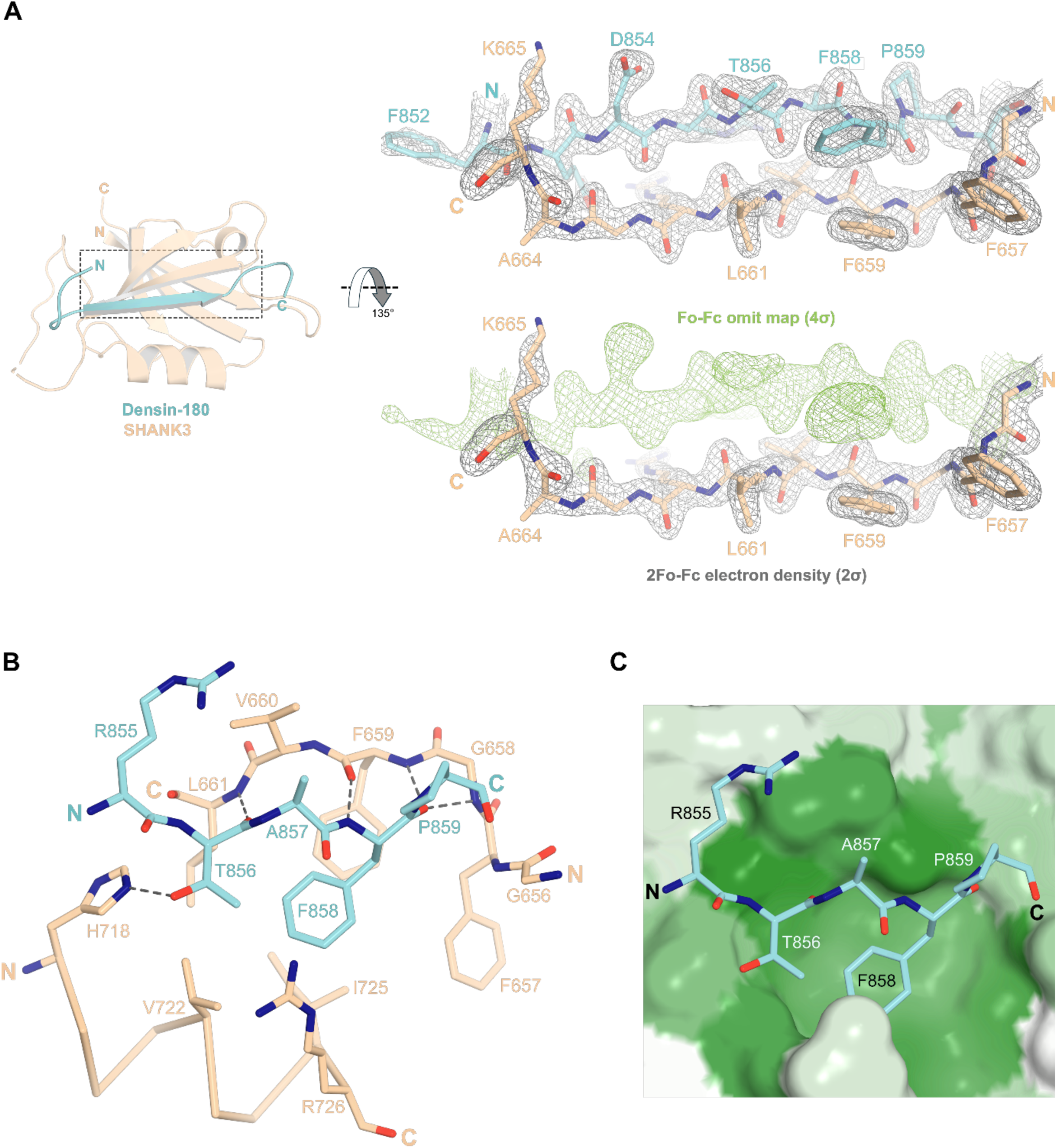
Crystal structure of the SHANK3 PDZ domain in complex with the internal Densin-180 PDZ-binding motif. (**A**) Left: overview of the SHANK3 PDZ domain in complex with the Densin-180 peptide (residues 843-864). SHANK3 is shown in wheat and Densin-180 in cyan. Right: 2Fo-Fc electron density map contoured at 2.0σ for the Densin-180 peptide (top) and simulated annealing Fo-Fc omit map contoured at 4.0σ (green mesh, bottom), confirming the position and conformation of the bound Densin-180 peptide. (**B**) Close-up view of the binding interface highlighting key interactions between Densin-180 and the SHANK3 PDZ domain, including the interaction between Thr856 and His718 and insertion of Phe858 into the hydrophobic pocket. Black dotted lines indicate hydrogen bonds. (**C**) Surface representation of the SHANK3 PDZ domain showing the Densin-180 peptide (residues 855-859) bound within the canonical ligand-binding groove. Phe858 occupies the conserved hydrophobic pocket, while Thr856 adopts the canonical class I PDZ ligand position. SHANK3 coloured by hydrophobicity using the AAindex scale FASG890101 (Nakai et al., 1988), where green indicates hydrophobic residues and white indicates polar residues.

